# The SAVED domain of the type III CRISPR protease CalpL is a ring nuclease

**DOI:** 10.1101/2024.05.08.593092

**Authors:** Sophie C. Binder, Niels Schneberger, Marianne Engeser, Matthias Geyer, Christophe Rouillon, Gregor Hagelueken

## Abstract

Prokaryotic CRISPR-Cas immune systems detect and cleave foreign nucleic acids. In type III CRISPR-Cas systems, the Cas10 subunit of the activated recognition complex synthesizes cyclic oligoadenylates (cOAs), second messengers that activate downstream ancillary effector proteins. Once the viral attack has been weathered, elimination of extant cOA is essential to limit the antiviral response and to allow cellular recovery. Various families of ring nucleases have been identified, specializing in the degradation of cOAs either as standalone enzymes or as domains of effector proteins. Here we describe the ring nuclease activity inherent in the SAVED domain of the cA_4_-activated CRISPR Lon protease CalpL. We characterize the kinetics of cA_4_ cleavage and identify key catalytic residues. We demonstrate that cA_4_-incuced oligomerization of CalpL is essential not only for activation of the protease, but is also required for nuclease activity. Further, the nuclease activity of CalpL poses a limitation to the protease reaction, indicating a mechanism for regulation of the CalpL/T/S signaling cascade. This work is the first demonstration of a catalytic SAVED domain and gives new insights into the dynamics of transcriptional adaption in CRISPR defense systems which are not aimed at abortive infection but rather at a reversible adaption to phage attack.

## Introduction

Bacteria are constantly threatened by the attack of foreign genetic elements, such as phages, and have evolved immune systems of varying complexity, such as restriction enzymes, toxin-antitoxin systems, CBASS, CRISPR, and many others as a countermeasure (1, 2). In CRISPR systems (3), a memory of previous phage attacks is stored in the form of short snippets of phage DNA in the CRISPR array of the bacterial chromosome (4). Those snippets are transcribed into short RNAs that are incorporated into recognition complexes sensing the presence of complementary foreign DNA or viral transcripts in the cell (5). If an attack is detected, the response can range from simple cleavage of the foreign DNA in the well-known Type II CRISPR systems (Cas9) (6) to complex multi-pronged responses in Type III CRISPR systems (7). In the latter, the Cas10 subunit of the recognition complex leads to the synthesis of cyclic oligoadenylates (cOAs), which are recognized by the CARF- or SAVED domains of effector proteins (8–10). The latter have a variety of biological functions such as DNAses, RNAses, nickases, transcription factors (7), or proteases as shown for CalpL (CRISPR-associated Lon protease), Craspase (CRISPR guided Caspase), the TPR-CHAT protease or the recently discovered SAVED-CHAT protein (10–13).

CalpL is part of the tripartite CalpL/T/S complex found in the thermophilic bacterium *Sulfurihydrogenibium* sp. YO3AOP1. The complex is formed by the protease CalpL, the anti-σ-factor CalpT and the ECF σ70 factor CalpS. Under normal conditions, the σ-factor is inhibited by CalpT. Under viral attack, cA_4_ is produced and the C-terminal SAVED domain of CalpL binds the second messenger with nanomolar affinity. This leads to oligomerization of CalpL and activation of the N-terminal Lon protease domain, which cleaves the 33 kDa anti-σ-factor CalpT, resulting in release of the 23 kDa CalpT_23_ fragment bound to the σ-factor CalpS. In analogy to other anti-σ-factor/σ-factor complexes, degradation by the ClpX/P-degron system is thought to release the sigma factor and allow the cell to adapt to viral attack (10).

Both bacteria and viruses have evolved mechanisms to either regulate or inhibit the antiviral response by degrading the cOA second messengers. This reaction is catalyzed by ring nucleases, which can be found as dedicated enzymes such as the archaeal host protein Sso2081 and the viral AcrIII-1 protein or as a second activity in the CARF domains of effector proteins such as the HEPN ribonuclease TTHB144 (7, 14–16).

Here, we show that the SAVED domain of the CalpL protease (Fig. 1 a) has ring nuclease activity and that the 2’-OH group of cA_4_ is very likely the nucleophile in the ring nuclease reaction (Fig. 1 b). We analyze the reaction by time-resolved HPLC/MS assays and study the influence of the nuclease activity on the oligomerization of CalpL and on its protease activity.

**Fig. 1.**
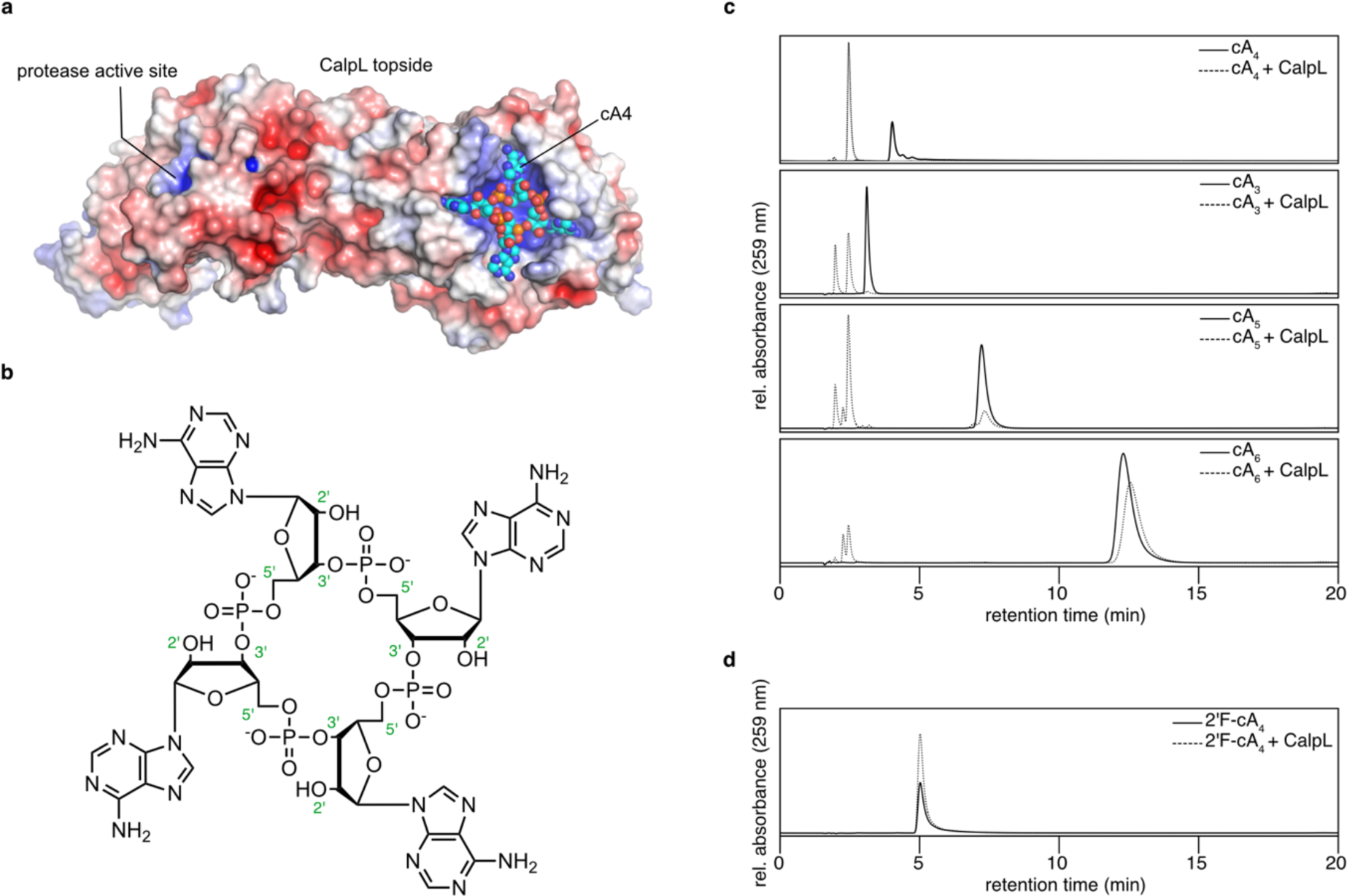
CalpL is a ring nuclease. **a)** A surface representation of a CalpL monomer with the electrostatic surface potential mapped onto the structure (red negative, blue positive). The cA_4_ molecule bound to the SAVED domain is shown in ball-and-sticks representation (PDB-ID: 8b0r, (10)). **b)** Structural formula of cA_4_. The numbering of selected atoms is given in green. **c, d)** HPLC traces recorded at 259 nm showing the result of incubating 30 µM of different cOAs with (dashed lines) and without (solid lines) 3 µM CalpL for 120 min. at 60 °C. HPLC traces are representative of at least three replicates.

## Results

### CalpL is a ring nuclease that degrades cA_4_ into two A_2_ units

To detect a possible ring nuclease activity of CalpL, we incubated CalpL with a 10-fold molar excess of cA_4_ and incubated the sample for 120 min at 60 °C, corresponding to the growth temperature of *Sulfurihydrogenibium* spp. An HPLC analysis of the sample clearly showed that the substrate peak had disappeared and a new peak with approximately 2-fold stronger intensity eluting at earlier retention times was observed (Fig. 1 c). This product peak was identified as linear di-AMP (A_2_) by comparison with HPLC standards and mass spectrometry (Fig. S1, Table S1).

In previous experiments, we identified cA_4_ as the activator of the Lon protease CalpL. SPR experiments revealed that CalpL binds cA_4_ with much higher affinity than other cOAs (10). Nevertheless, to a small extent, the protease activity of CalpL was also stimulated by other cOAs (cA_3_, cA_5_, cA_6_). While this activation by other cOAs is very likely not of any physiological relevance, it is interesting from a mechanistic point of view, because the activation mechanism appears to be “flexible” enough to accommodate the different sizes of the cOAs. We thus wondered whether the CalpL ring nuclease activity is specific towards cA_4_ and performed nuclease assays using different cOAs (cA_3_, cA_5_, cA_6_) as substrates (Fig. 1 c). Interestingly, we observed significant cleavage of cA_3_ and cA_5_ within 120 min at 60 °C, whereas cA_6_ was degraded only to a minor extent. For cA_3_, we observed an almost complete conversion into two cleavage products, which were identified as 2’,3’-cAMP and A_2_ by comparison with HPLC standards. For cA_5_, we observed an incomplete cleavage of cA_5_ to A_2_>P (5’-ApAp with a cyclic 2’,3’ phosphate), A_2_ and 2’,3’-cAMP. For cA_6_, we observed traces of A_2_>P, A_2_ and 2’,3’-cAMP, however most of the substrate remained uncleaved.

By analogy with other nucleases and ring nucleases, we wondered, whether the 2’-OH group of the cOA might be the nucleophile of the reaction. To test this, we performed the same experiment with the cA_4_ derivative 2’F-cA_4_ containing fluoro modifications in all ribose 2’ positions (Fig. 1 d). In this case, no substrate cleavage was observed within 120 min at 60 °C, suggesting that the nuclease reaction indeed proceeds via positioning of the 2’-OH of the ribose for the nucleophilic attack.

To assess the impact of CalpL nuclease activity on the CalpL/T/S cascade, we tested the cA_4_ degradation products for activation of the CalpL protease activity. As expected, we observed a strong proteolytic cleavage of CalpT in the presence of cA_4_ within 120 min at 60 °C (Fig. 2 a). Substitution of cA_4_ with A_4_ (with a 3’ phosphate) led to a strongly decreased but still detectable amount of CalpT cleavage as judged by Coomassie-stained SDS-PAGE. For A_2_ (with a 3’ phosphate), we did not observe CalpT cleavage, demonstrating that cA_4_ degradation by CalpL limits the protease activity.

**Fig. 2.**
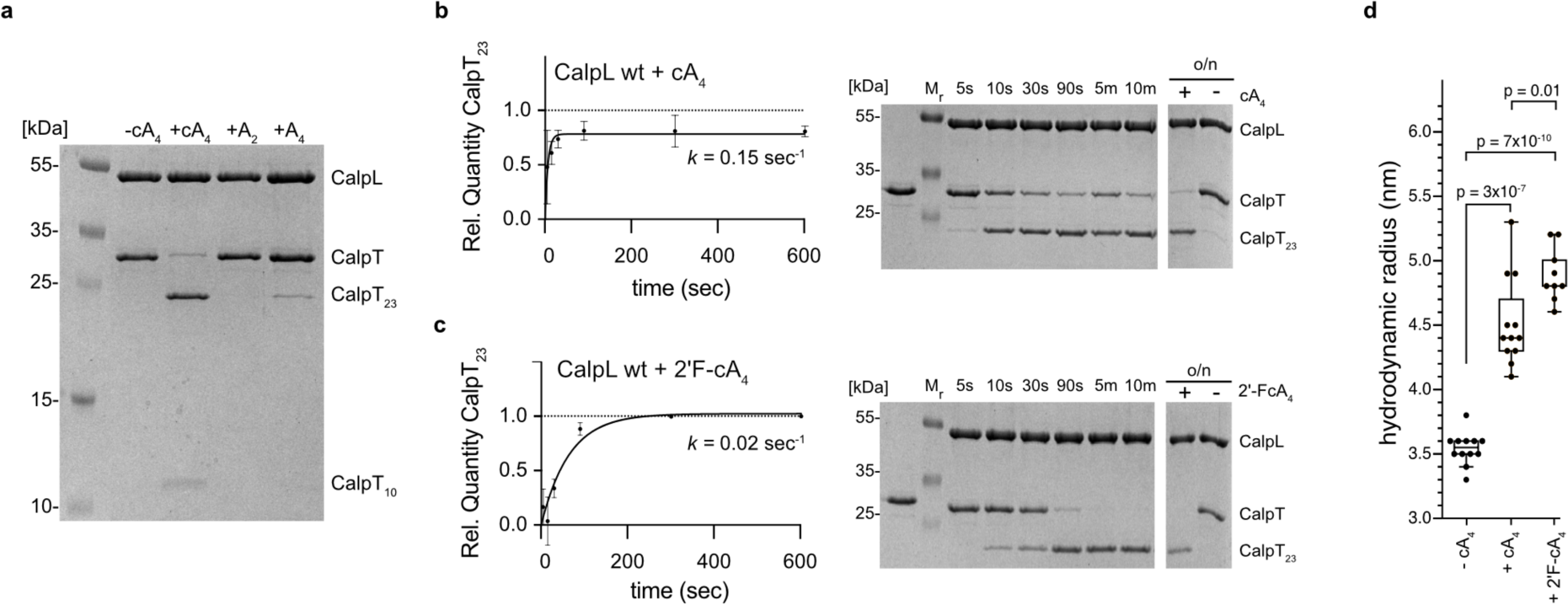
Degradation of cA_4_ limits the protease activity. **a)** Coomassie-stained SDS-PAGE analysis of CalpL protease activity assay showing the amount of CalpT cleavage upon addition of cA4, A_2_, or A_4_. For the assay, 3 µM of CalpL and CalpT were mixed with 3 µM of the respective adenylate compound and incubated for 120 min. at 60 °C. **b)** Quantification of CalpT cleavage at 37 °C for the indicated time periods of incubation of CalpL and CalpT with cA_4_ as described for a). Right: Example of Coomassie-stained 15 % SDS-PAGE gel used for quantification. All experiments were performed in triplicates and error bars represent mean ± SD. **c)** same as in b) but in the presence of 2’F-cA_4_. **d)** Hydrodynamic radii of CalpL determined at a sample concentration of 86 µM (5 mg/ml) in the presence or absence of cA_4_ or 2’F-cA_4_. All DLS experiments were performed in triplicates. Bars display median and interquartile range; *p*-values for two-tailed t-tests are indicated.

Based on the observation that the protease activity of CalpL is strongly reduced upon linearization of cA_4_, we speculated whether 2’F-cA_4_ might further enhance the proteolytic activity of CalpL. To test this, we performed protease reaction time courses by incubating equimolar concentrations of CalpL and CalpT in the presence of either cA_4_ or 2’F-cA_4_ (Fig. 2 b, c). Further, we studied the oligomerization dynamics of CalpL in the presence of cA_4_ or 2’F-cA_4_ by determining the hydrodynamic radii using dynamic light scattering (DLS) (Fig. 2 d). Indeed, DLS analysis revealed a slightly increased hydrodynamic radius for CalpL of 4.9 ± 0.21 nm upon binding of 2’F-cA_4_, compared to 4.5 ± 0.35 nm upon addition of cA_4_. Despite a slower initial velocity (v_0_) of the reaction, we observed complete consumption of CalpT after 5 minutes of incubation with 2’F-cA_4_ at 37°C, whereas activation with cA_4_ failed to induce full cleavage of CalpT even after overnight incubation.

### Nuclease reaction time courses and identification of intermediates

To follow the reaction resulting in the conversion of cA_4_ to A_2_, we performed time course experiments and analyzed the reaction products by HPLC/MS (Fig. 3, Supplementary Table S1). Due to the high speed of the ring nuclease reaction, we lowered the reaction temperature from 60 °C to 37 °C, resulting in a reduced activity of CalpL. The reactions were stopped by flash freezing in liquid nitrogen and thawed immediately before injection onto the column. Five seconds after the start of the reaction, two additional peaks with higher retention times appeared. Using MS, these were identified as A_4_ (larger peak, 4.4 min.) and A_4_>P (smaller peak, 4.1 min.). Within 30 seconds of incubation, a large fraction of cA_4_ was consumed and converted into A_4_, with only a minor A_4_>P fraction being present at all time points tested. In addition, two smaller peaks at much earlier retention times had appeared. MS analysis revealed that the peaks had the mass of linear A_2_ (2.4 min.) and of A_2_>P (2.2 min.). All assignments were confirmed by comparison with HPLC standards (Fig. S 2). Over the next minutes, the A_2_>P intermediate was almost completely converted to A_2_. Note that we observed very small amounts of A_1_>P (1.9 min.), but no A_3_>P, being formed after extend incubation periods. In summary, the CalpL ring nuclease reaction proceeds from cA_4_ via A_4_>P, A_4_, A_2_>P to A_2_.

**Fig. 3.**
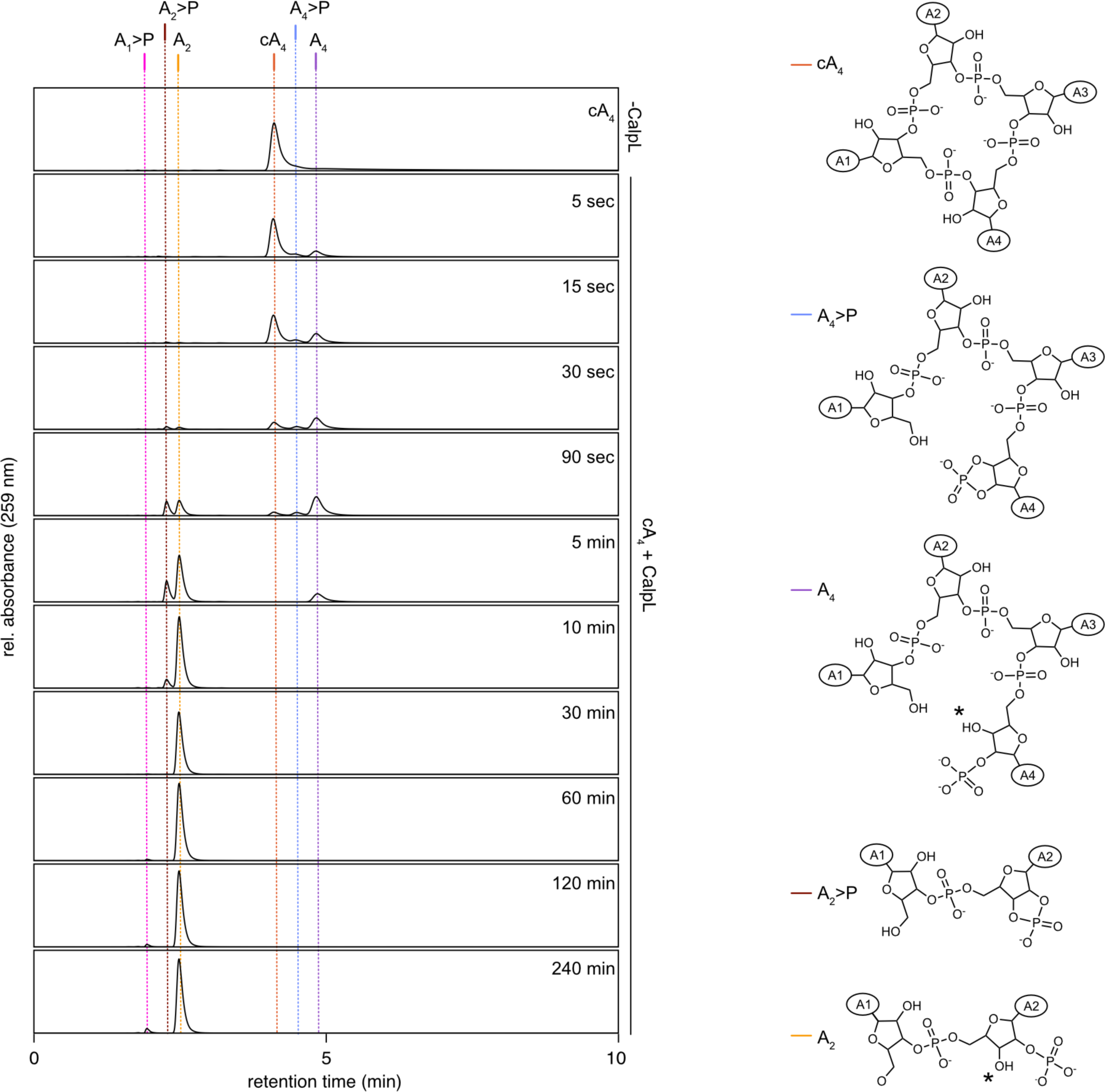
Time course of CalpL-mediated cA_4_ cleavage. Left: HPLC traces of nuclease reactions incubating 1.5 µM CalpL with 15 µM cA_4_ at 37 °C for the indicated time periods. The vertical lines mark reaction intermediates and products that were identified by comparison with standard runs and/or mass spectrometry. Right: Structural formulas of the observed reaction intermediates. The adenosine base is represented by A_1_-A_4_ in ellipsoids. All HPLC traces are representative of at least three replicates. The asterisk symbol (*) indicates that hydrolysis of the cyclic 2’,3’-phosphate could yield products having the phosphate group at either the 2’ or 3’ position.

### Identification of residues involved in ring nuclease activity of CalpL

Considering the fact that cA_4_ is split into two A_2_ molecules, we thought it likely that the cleavage takes place at two opposing positions in the active site. Based on structural data obtained from the CalpL-cA_4_ complex (Fig. 1 a) combined with sequence alignments of CalpL homologs, we identified conserved side chains, which are located in the cA_4_ binding site and could be involved in the nuclease reaction: H345, H392, H474, S325, S391, S451 (Fig. 4 a).

**Fig. 4.**
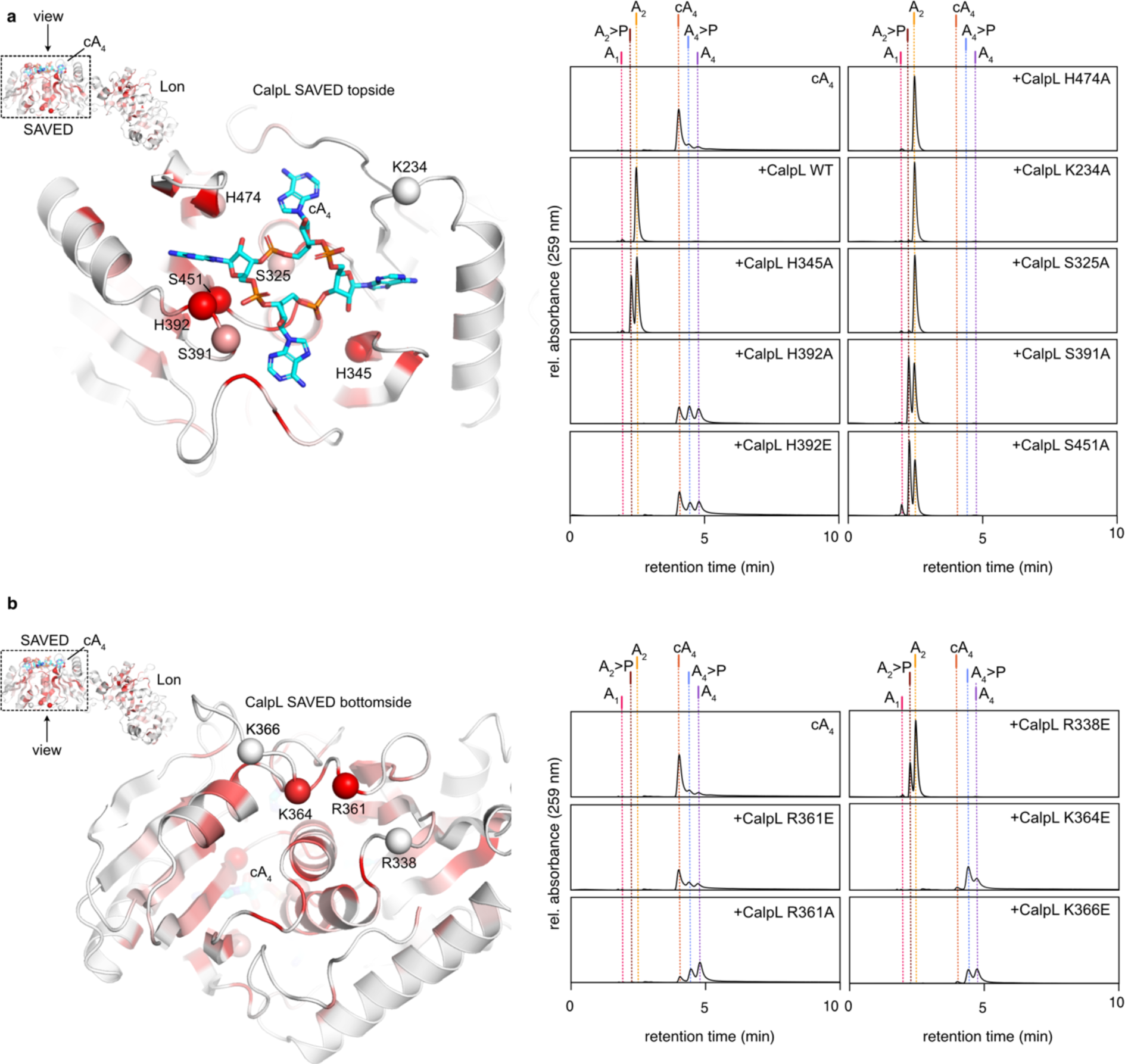
CalpL mutants in the cA_4_ binding site and on the bottom of the SAVED domain show impaired ring nuclease activity. **a)** Left: A cartoon model of the SAVED domain of CalpL (viewing direction with respect to the complete structure is indicated). The conservation of residues is mapped onto the structure (red: high conservation score, white low conservation score; The underlying alignment is provided as Supplementary Material). The cA_4_ molecule is shown in cyan sticks. Residues that were mutated are marked by a sphere at their Cα position. Right: HPLC analysis of ring nuclease reactions incubating 1.5 µM of the respective SAVED topside variant indicated in a) with 15 µM cA_4_ at 60 °C for 120 min. **b)** Left: As in a) but the bottom side of the SAVED domain is shown. Right: HPLC analysis of ring nuclease reactions performed as in a) but of SAVED bottom side indicated in b). All HPLC traces are representatives of at least three replicates.

To assess their contribution to the ring nuclease reaction, we mutated the conserved residues to alanine, either individually or in combination, and examined the cA_4_ cleavage reaction at 60 °C by HPLC analysis.

The strongest effect of nuclease attenuation was observed for mutants H392A and for H392E. Here we observed a slow conversion of cA_4_ to A_4_>P and A_4_, but we did not observe any production of di-adenylates after incubation for 120 min. Whereas mutation of either S325 or H474 to alanine did not affect the nuclease activity, H345A and S391A variants of CalpL showed a reduced ability to convert A_2_>P to A_2_ within the observed time frame. We suspected a potential cooperation of H345 and H392, however surprisingly, a combined mutation to alanine resulted in an enhanced nuclease activity compared to mutation of H392 alone (Fig. S3). Here, we observed production of A_2_>P and A_2_, with only a minor fraction of A_4_ present after incubation at 60 °C for 120 min. Interestingly, we further observed production of a novel reaction product which was identified as A_3_>P by comparison with HPLC standards (Fig. S3). Although not conserved, we considered K234, located directly opposed to H392, as a possible candidate for a coordinated cA_4_ cleavage. As judged by the reaction endpoint analysis, mutation of K234 to alanine did not influence the nuclease activity of CalpL. Similarly to the H345A/H392A variant, a combined mutation of K234 and H392 to alanine enhanced the nuclease activity compared to the H392 single alanine mutant, and further resulted in the production of minor traces of A_3_>P. A combined alanine mutation of H345 and K234 did not enhance the nuclease attenuating effect observed for H345A alone and did not show production of tri-adenylate species.

In summary, of all residues inside the cA_4_ binding site, only residues in the vicinity of H392 showed a strong impact on the ring-nuclease activity, suggesting that at least one cleavage reaction occurs at this position.

Puzzled by the persistence of nuclease activity despite systematic mutation of residues within the cA_4_ binding pocket, our attention was drawn to the high conservation of a positively charged patch on the backside of the SAVED domain (Fig. 4 b). Recent reports on CARF domain-containing ring nucleases and oligomerizing SAVED domains (17, 18) prompted us to investigate whether conserved bottom side residues might be involved in the ring nuclease reaction when CalpL forms oligomers. We mutated the corresponding residues, either to alanine or glutamate, and assessed the reaction products via HPLC. Strikingly, all backside mutants showed a significantly impaired nuclease activity, with none achieving a complete conversion of cA_4_ to A_2_ within 120 min at 60°C. While K364E and K366E variants showed a minimal production of A_4_>P and A_4_ within the observed time frame, mutation of R361 to glutamate resulted in a complete disruption of the nuclease activity. Interestingly, an R361A variant retained basal levels of nuclease activity and showed production of A_4_>P and A_4_.

These observations indicate a strong link between oligomerization and nuclease activity; however, the exact allocation of active site residues is not trivial.

### Interdependence of the nuclease and protease activities of CalpL

Fig. 5 a shows a model of a CalpL oligomer based on the stacked SAVED domains of the recently published SAVED-CHAT structure(11). It illustrates that conserved residues of both, the CalpL bottom- and top sides, including R338, R361, K364, and K366 are in close contact to the sandwiched cA_4_ molecule, rationalizing their influence on the ring nuclease activity.

**Fig. 5.**
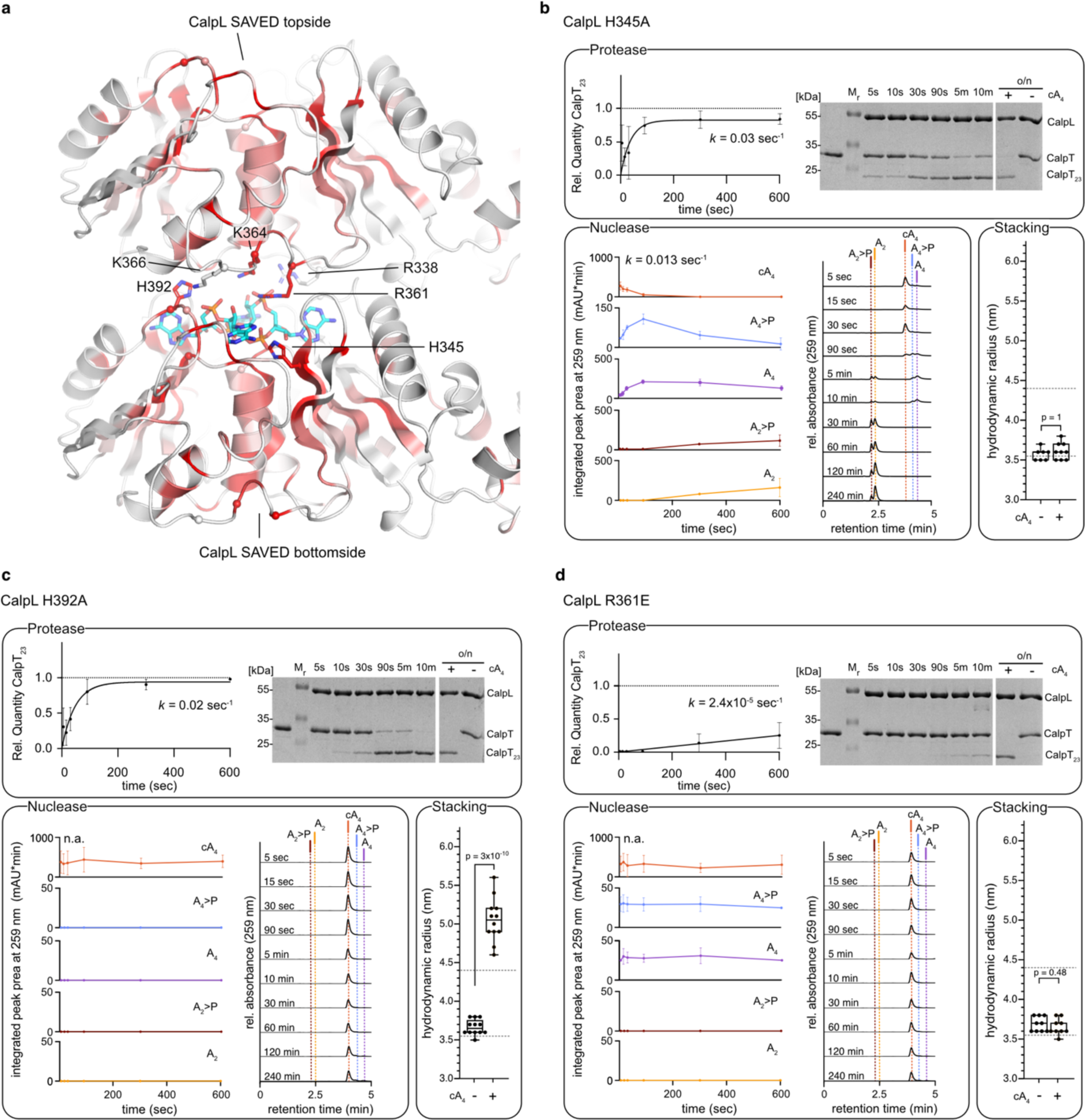
Interdependence of ring nuclease and protease activities of CalpL. **a)** An energy-minimized model for cA_4_-mediated oligomerization of CalpL based on the SAVED-CHAT structure (PDB-ID: 8tl0, (11)) generated with GROMACS (19). The cA_4_ molecule is shown as cyan sticks. **b-d)** Upper panels: Quantification of CalpT cleavage by incubating 3 µM of the respective CalpL mutant with 3 µM CalpT at 37 °C for the indicated time points. A Coomassie-stained SDS-PAGE gel used for quantification is included for reference. Left lower panels: Quantification of nuclease reaction intermediates and products generated upon incubation of 1.5 µM CalpL and 15 µM cA_4_ at 37 °C for the indicated time points. HPLC traces used for quantification by peak area integration at 259 nm are shown exemplary. Right lower panels: Hydrodynamic radii of CalpL determined at a sample concentration of 86 µM (5 mg/ml) in the presence or absence of cA_4_. Dashed lines mark the hydrodynamic radii of wildtype CalpL with (4.5 ± 0.35 nm) and without (3.5± 0.12 nm) cA_4_. All experiments were performed in triplicates. Bars of DLS data display median and interquartile range; *p*-values for two-tailed t-tests are indicated.

Activation of the ring nuclease upon oligomer formation and protease activation constitutes an elegant mechanism to limit the protease activity. Consequently, we wondered if and how a reduced nuclease activity, such as observed for top side residue H392A or bottom side residues R361A/E, K364E, and K366E might influence the protease activity of CalpL.

To study this, we performed protease time course experiments incubating the different CalpL variants and the protease substrate CalpT and quantified the protease activities using Coomassie-stained SDS-PAGE (Fig. 5b-d, Fig. S5). For each experiment we assessed the formation of CalpL oligomers by determining hydrodynamic radii using dynamic light scattering (DLS) (Fig. 5 b-d, Fig. S4).

DLS analysis revealed a strongly enhanced cA_4_-induced oligomerization for the nuclease-deficient H392A topside mutant as judged by a hydrodynamic radius of 5.1± 0.28 nm, similar to the hydrodynamic radius of 4.9± 0.21 nm observed upon 2’F-cA_4_-induced oligomerization of the wildtype protein shown in Fig. 2. Analogous to the activation of wildtype CalpL with 2’F-cA_4_, the protease reaction of the H392A variant showed a reduced initial velocity V_0_ but achieved full cleavage of CalpT within 10 min (*k* [protease] = 2.7×10^-4^ sec^-1^).

The CalpL H345A variant, which had an impaired capacity to convert A_2_>P to A_2_ in the nuclease reaction endpoint experiment, showed similar kinetics of cA_4_ cleavage in the time-course experiment (*k* [H345A; nuclease] = 0.013 sec^-1^; *k* [wt; nuclease] = 0.017 sec^-1^) (Fig. S5). Interestingly, however, mutation of H345A resulted in a decreased protease activity (*k* [H345A; protease] = 0.03 sec^-1^; *k* [wt; protease] = 0.15 sec^-1^) with a minor fraction of uncleaved CalpT after incubation overnight. Further, we did not observe oligomerization of H345A in response to cA_4_, as evidenced by a similar hydrodynamic radius in the presence (3.6± 0.11 nm) and absence (3.6± 0.13 nm) of cA_4_. Intriguingly, for the H474A variant, we observed a strong ring nuclease activity (*k* [H474A; nuclease] = 0.058 sec^-1^) (Fig. S5), which, in contrast to the wildtype protein, did not correlate with an early termination of the protease reaction. Here, we observed a complete substrate cleavage within 10 min despite an almost complete consumption of A_4_ after 5 min. The oligomerization behavior, however, remained unaltered as judged by a hydrodynamic radius of 3.6 ± 0.11 nm in the absence and 4.3 ± 0.22 nm in the presence of cA_4_ (Fig. S5).

In the case of the nuclease-dead CalpL variant R361E, we observed a strong decrease in protease activity (*k* [R361E; protease] = 2.4×10^-5^ sec^-1^), correlating with the absence of cA_4_-induced oligomerization, as indicated by a similar hydrodynamic radius both in the absence (3.7± 0.09 nm) and presence (3.7± 0.10 nm) of cA_4_. Conversely, CalpL R361A, which showed basal levels of nuclease activity, exhibited strong proteolytic activity (*k* [R361A; protease] = 0.03 sec^-1^), while showing only a marginal cA_4_-induced oligomerization, as judged by a hydrodynamic radius of 3.8± 0.12 nm (Fig. S5). It should be kept in mind our DLS experiments are not sensitive enough to capture sporadic oligomerization events and our results indicate that even such transient cA_4_-induced oligomerization of CalpL appears to be sufficient to trigger cleavage of CalpT.

## Discussion

Here we have characterized the ring nuclease activity of the CalpL SAVED domain. The protein rapidly degrades its activator, the cyclic oligonucleotide cA_4_ to linear A_2_ via several intermediates with cyclic phosphate groups. We did not observe any indications of a metal dependency of the reaction and the fact that the chemical analog 2’F-cA_4_ was not degraded, together with the observation of cyclic phosphate intermediates are strong indicators of the 2’-OH group of the cA_4_ molecule being the nucleophile of the reaction.

Our previously determined structure of CalpL in complex with cA_4_ allows us to speculate about the location of the scissile bonds in the cA_4_ molecule with respect to the surrounding protein surface. We found a strong effect on the rate of the A_2_>P to A_2_ reaction when residues near H392 were mutated. This indicates that at least one cleavage reaction takes place at this site. Since the final product is linear A_2_, a second cleavage must occur and the logical position for this reaction would be exactly opposite of H392. However, we did not observe strong effects on the nuclease reaction when residues at this position in the cA_4_ binding site were mutated, such as K234A. In principle, one could imagine that the cA_4_ molecule dissociates and reassociates with the SAVED domain between two cleavage reactions. However, in this case we would have expected to observe significant production of A_1_>P/ A_1_ and A_3_>P/A_3_, which was clearly not the case. Instead, our data indicates that residues from the bottom side of CalpL oligomers are involved in the formation of the ring nuclease active site, although it is currently unclear whether they play a catalytic or merely a structural role.

We found that the ring nuclease activity of CalpL is severely decreased for mutants that also show a decreased capability to form CalpL oligomers, such as H345A, R361A/E, K364E, and K366E. This would ensure that the ring nuclease activity is only fully activated when the cA_4_ concentration is high enough to allow the formation of oligomers and would thereby avoid a decrease in sensitivity by premature hydrolysis of cA_4_ molecules. Since even the mutants might undergo sporadic oligomerization, we conclude that the remaining nuclease activity of these mutants is due to such rare events. A high-resolution structure of a stacked SAVED domains of CalpL would certainly be helpful to further unravel the mechanism of the ring nuclease reaction. In previous SAXS experiments (10), we observed indications of a staggered arrangement of CalpL monomers in solution, though at low resolution. This is quite different to the non-staggered arrangement found in the SAVED-CHAT cryo-EM structure (11). Our model based on the latter structure can rationalize the effects of bottom-side mutants analyzed in this study very well. Since the cA_4_ binding site is pseudo C2 symmetric, it might be that both arrangements can occur in solution, especially considering the fact that CalpL is normally tightly bound to the CalpT and CalpS proteins.

Studies exploring the molecular mechanisms of type III CRISPR associated ring nucleases have described a diverse array of nuclease activation processes. These mechanisms encompass intricate conformational changes within the cOA binding site, leading to an alignment of the active site (20, 21), as well as global structural rearrangements including multimerization, where individual nucleases assemble to form a composite active site from previously isolated regions of the protein (22). CalpL is the first protein with a SAVED domain to show a ring nuclease activity and appears to be a mix of features observed in other ring nucleases.

Most likely, the CalpL nuclease activity will lead to a self-quenching effect of the antiviral response, as observed for other type III CRISPR systems. This indicates that the CalpL/T/S pathway is not aimed at an abortive infection but rather at a reversible adjustment of the organism to a phage attack where even a small amount of released CalpS might be enough to appropriately fine tune the transcriptional levels of the cell. Alternatively, the ring nuclease activity might also play a role in setting up an activation threshold such that basal levels of cA_4_ are quickly degraded. However, since the nuclease activity appears to go hand-in-hand with oligomer formation and thereby the activation of the protease activity of CalpL the self-quenching effect appears to be the more likely explanation for its ring nuclease activity.

The CalpL protein combines a staggering number of functionalities in a relatively small protein. The inducible protease activity makes it a potent tool for biotechnological applications, such as for instance for CRISPR-based sensing assays, where the protease activates a response mechanism. Due to its negative impact on the protease activity, the ring nuclease activity is a disadvantage for such applications. Therefore, our discovery that it can be deactivated while maintaining the protease activity is of high interest for such efforts.

## Materials and methods

### Generation of CalpL expression vectors

All CalpL constructs were expressed from a pET11a plasmid containing the codon-optimized sequence with an N-terminal 10xHis-TEV tag. Point mutations were introduced into plasmids by primer-directed mutagenesis. The non-mutated parental plasmid was digested by the methylation-sensitive restriction endonuclease *Dpn*I. After restriction digestion, 10 µl of the reaction mixture was used for transformation of NEB DH5α cells.

### Expression and purification of CalpL

Recombinant His10-CalpL was expressed in *Escherichia coli* BL21 (DE3) bacterial cells. *E. coli* cells were grown in LB medium containing the appropriate antibiotic at 37 °C for 16 h (pre-culture). The following day, larger volumes of LB medium were inoculated with the pre-culture and adjusted to an optical density (OD_600_) to 0.1. Cultures were grown at 37 °C to an OD_600_ of 0.8 and protein expression was induced by addition of IPTG to a final concentration of 0.4 mM. Proteins were expressed at 20 °C for 16h. Bacteria were harvested by centrifugation at 4,000 x*g* for 25 min. Cell pellets were snap-frozen in liquid nitrogen and stored at −20 °C or subjected to immediate cell lysis.

Cell pellets were resuspended in lysis buffer (20 mM Tris-HCl pH 8.0, 50 mM NaCl) and lysed by sonication. Cell debris was sedimented by centrifugation at 25,000 x*g* for 45 min at 10 °C. The lysate was filtered through a membrane filter with a 0.8 µm pore size and subjected to affinity chromatography.

For affinity chromatography, Ni^2+^-NTA resin was equilibrated with lysis buffer and incubated with lysate for 2 h at 4 °C on a rolling incubator. The resin was then transferred to a gravity column and washed extensively with wash buffer (20 mM Tris-HCl pH 8.0, 50 mM NaCl, 20 mM imidazole) and eluted with elution buffer (20 mM Tris-HCl pH 8.0, 50 mM NaCl, 500 mM imidazole). The elution fraction was dialyzed against lysis buffer for 16 h at 4 °C and subjected to further purification by Heparin chromatography.

Heparin chromatography was performed on a HiPrep Heparin Fast Flow 16/60 column (Cytiva) equilibrated in binding buffer (20 mM Tris-HCl pH 8.0) using an ÄKTA FPLC system (Cytiva). The column was washed extensively with binding buffer and proteins were eluted by increasing the concentration of NaCl from 0 mM to 1 M over 3.6 column volumes (CV). Fractions containing large amounts of CalpL were pooled and digested with TEV protease (1:25 *w/w*) for 16 h at 4 °C for removal of the 10xHis-tag.

Following protease digest, CalpL was further purified by size exclusion chromatography (SEC) on a HiLoad 16/600 Superdex 75 pg column (Cytiva) equilibrated with lysis buffer and reverse Ni^2+^-NTA purification for removal of residual tag, non-cleaved protein and TEV protease.

### Expression and purification of CalpT

CalpT was expressed from a pBAD plasmid containing the codon-optimized sequence with an N-terminal 6xHis-TEV tag. Recombinant 6xHis-CalpT was expressed in *Escherichia coli* BL21 (DE3) bacterial cells. *E. coli* cells were grown in LB medium containing the appropriate antibiotic at 37 °C for 16 h (pre-culture). The following day, larger volumes of LB medium were inoculated with the pre-culture and adjusted to an optical density (OD_600_) of 0.1. Cultures were grown at 37 °C to an OD_600_ of 0.8 and protein expression was induced by addition of 0.5 g/L L(+)-arabinose. Proteins were expressed at 30 °C for 5 h. Bacteria were harvested by centrifugation at 4,000 x*g* for 25 min. Cell pellets were snap-frozen in liquid nitrogen and stored at −20 °C or subjected to immediate cell lysis.

Cell pellets were resuspended in lysis buffer (25 mM Tris-HCl pH 8.0, 500 mM NaCl, 1 mM DTT, 10 % glycerol) and lysed by sonication. Cell debris was sedimented by centrifugation at 25,000 x*g* for 45 min at 20 °C. The lysate was filtered through a membrane filter with a 0.8 µm pore size and subjected to affinity chromatography.

For affinity chromatography, Ni^2+^-NTA resin was equilibrated with lysis buffer and incubated with lysate for 2 h at room temperature on a rolling incubator. The resin was then transferred to a gravity column and washed extensively with wash buffer (25 mM Tris-HCl pH 8.0, 500 mM NaCl, 1 mM DTT, 10 % glycerol, 40 mM imidazole) and eluted with elution buffer (25 mM Tris-HCl pH 8.0, 500 mM NaCl, 1 mM DTT, 10 % glycerol, 1 M imidazole). The elution fraction was dialyzed against binding buffer (25 mM Tris-HCl pH 8.0, 1 mM DTT, 10 % glycerol) for 16 h at room temperature and subjected to further purification by Heparin chromatography.

Heparin chromatography was performed on a HiPrep Heparin Fast Flow 16/60 column (Cytiva) equilibrated in binding buffer using an ÄKTA FPLC system (Cytiva). The column was washed extensively with binding buffer and proteins were eluted by increasing the concentration of NaCl from 0 mM to 500 mM over 3.6 CVs. Fractions containing large amounts of 6xHis-CalpT were pooled and further purified by SEC on a HiLoad 16/600 Superdex75 pg column (Cytiva) equilibrated with 25 mM Tris-HCl, 500 mM NaCl, 1 mM DTT, 10 % glycerol.

### Nuclease assay and RP-HPLC analyses

For testing the nuclease activity, CalpL was incubated with 10-fold molar excess of the respective cOA in 20 mM Tris-HCl, 50 mM NaCl at either 37 °C (time-course measurement) or 60 °C (endpoint measurement) and analyzed by reversed-phase high-performance liquid chromatography (RP-HPLC) using the Infinity II HPLC system (Agilent). Nuclease reaction products were separated at a flow-rate of 1 ml*min^-1^ on a Chromolith Performance RP-18e 100 x 46 mm column (Merck) equipped with a guard cartridge. The eluent was composed of 30 mM K_2_HPO_4_, 70 mM KH_2_PO_4_, 10 mM tetrabutylammonium bromide, 13 % acetonitrile (*w/w*). RNA species were collected manually and subjected to mass spectrometry for identification.

For calculation of rate constants *k*, the fraction of cA_4_ cut was determined by integrating the peak area at 259 nm for the intervals of t = 5 sec. to t = 600 sec. Values were normalized to t = 5 sec. For the CalpL variants H345A, H474A, and wildtype, inversed values were fitted by a one phase decay model (Y=(Y0-Plateau)*exp(-K*X)+Plateau) to the time points intervals of t = 5sec to t = 600 sec using Prism 9.5.0. Due to the slow progression of the nuclease reaction catalyzed by the CalpL variants H392A, R361A, and R361E, no adequate fit was applicable.

### Generation of HPLC standards

For generation of A_4_>P, A_3_>P, and A_2_>P as HPLC standards similar to (23), 30 µM of MazF substrate (for A_4_>P: 5’-*aaaa*acacugaaccug-3’; for A_3_>P: 5’-*aaa*acacugaaccug-3’ for A_2_>P: 5’-*aa*acacugaaccug-3’) were incubated with 10 U recombinant MazF (Takara Bio) in 20 mM Tris pH 8.0, 50 mM NaCl and incubated for 60 min @ 37 °C. Linear adenylate standards were ordered containing a 3’phosphate modification. MazF substrates and linear oligoadenylates were supplied by Metabion (Germany).

### Mass spectrometry analyses

Samples manually collected after HPLC separation were subjected to HPLC-MS analysis with a Q/TOF mass spectrometer (Bruker micrOTOF-Q) equipped with an electrospray ion source and a HPLC system (Agilent 1200 series) using a RP 150 x 2 mm C18 column (Knauer Eurospher II 100-5C18). The flow rate was set to 0.25 ml*min^-1^ with a linear gradient starting from a 95:5 mixture of 25 mM aqueous ammonium acetate and acetonitrile ramping to a 30:70 ratio after 15 minutes. The column was rinsed with 5:95 NH_4_OAc:acetonitrile after every run. RNA samples were detected via UV absorption at 254 nm and ESI spectra measured in negative mode.

### Protease assay

To monitor protease activity over time, CalpL and 6xHis-CalpT were combined at 3 µM each in 20 mM Tris-HCl pH 8.0, 50 mM NaCl, and incubated at 37 °C for 30 min. For activation of the protease reaction, cA_4_ was added to a final concentration of 3 µM, and the reaction proceeded at 37 °C for the indicated time periods. For endpoint analysis of protease reactions, the same protocol was followed but with a 30-minute incubation at 60 °C. After addition of cyclic or linear adenylate species, the reactions were incubated for further 60 minutes at 60 °C. Reactions were stopped by addition of reducing SDS loading buffer and heating for 10 min at 94 °C. The samples were loaded on a 15 % SDS polyacrylamide gel, run at 40 mA/gel for 60 min and analyzed by Coomassie staining.

Cleavage of 6xHis-CalpT was assessed by quantifying CalpT bands on Coomassie-stained SDS-PAGE gels and normalizing to a non-cleaved 6xHis-CalpT control sample using the Image Lab Software (BioRad). Inverse values were calculated for plotting. Due to an inadequate integration of CalpT band intensities for the CalpL R361E variant, quantification was done by integrating band intensities of CalpT_23_ and normalization to the amount of CalpT_23_ at t = 240 min. Rate constants *k* were calculated by fitting a one phase decay model (Y=(Y0-Plateau)*exp(-K*X)+Plateau) to the time points intervals of t = 5sec to t = 600 sec using Prism 9.5.0.

### Dynamic light scattering

To investigate cA_4_ induced oligomerization, the hydrodynamic radii of CalpL in the presence or absence of cA_4_ or 2’F-cA_4_ were measured by dynamic light scattering using the DynaPro NanoStar system (Wyatt Technology). Samples containing CalpL at a final concentration of 86.6 µM (5 mg/ml) were prepared in 20 mM Tris-HCl pH 8.0, 50 mM NaCl and centrifuged at 15,000 x*g* for 15 min to remove aggregates before each measurement. For each condition, three measurements were performed at a sample temperature of 20 °C and three measurement cycles of each 20 single data acquisitions with acquisition times of 3 sec. To investigate the significance of the radii difference between samples, a two-tailed unpaired t-test was performed (*p* values are included in the corresponding figures).

### Structural modelling

To construct a model of a cA_4_ induced oligomer of CalpL, two individual cA_4_ bound CalpL monomers (PDB-ID: 8b0r) were superimposed on the dimeric SAVED-CHAT structure (PDB-ID: 8tl0) with PyMOL (www.pymol.org). This led to a model with some clashes between the two protomers. The CHARMM-GUI (24) and GROMACS (19) were then used to perform an energy minimization of the model, including the sandwiched cA_4_ molecule.

## Acknowledgements

We thank Malcolm White and Haotian Chi (University of St Andrews) for discussions. GH is grateful for funding by the German Research Foundation (DFG, grant number HA-6805/6-1).

## Author contributions

SB performed experiments, analyzed data and wrote the manuscript together with GH. NS performed experiments and analyzed data. ME performed experiments and analyzed data. MG analyzed data. CR performed initial experiments with GH and analyzed data. GH analyzed data, supervised the study and acquired funding. All authors contributed to the writing of the manuscript.

## Supplementary Information

**Fig. S1.**
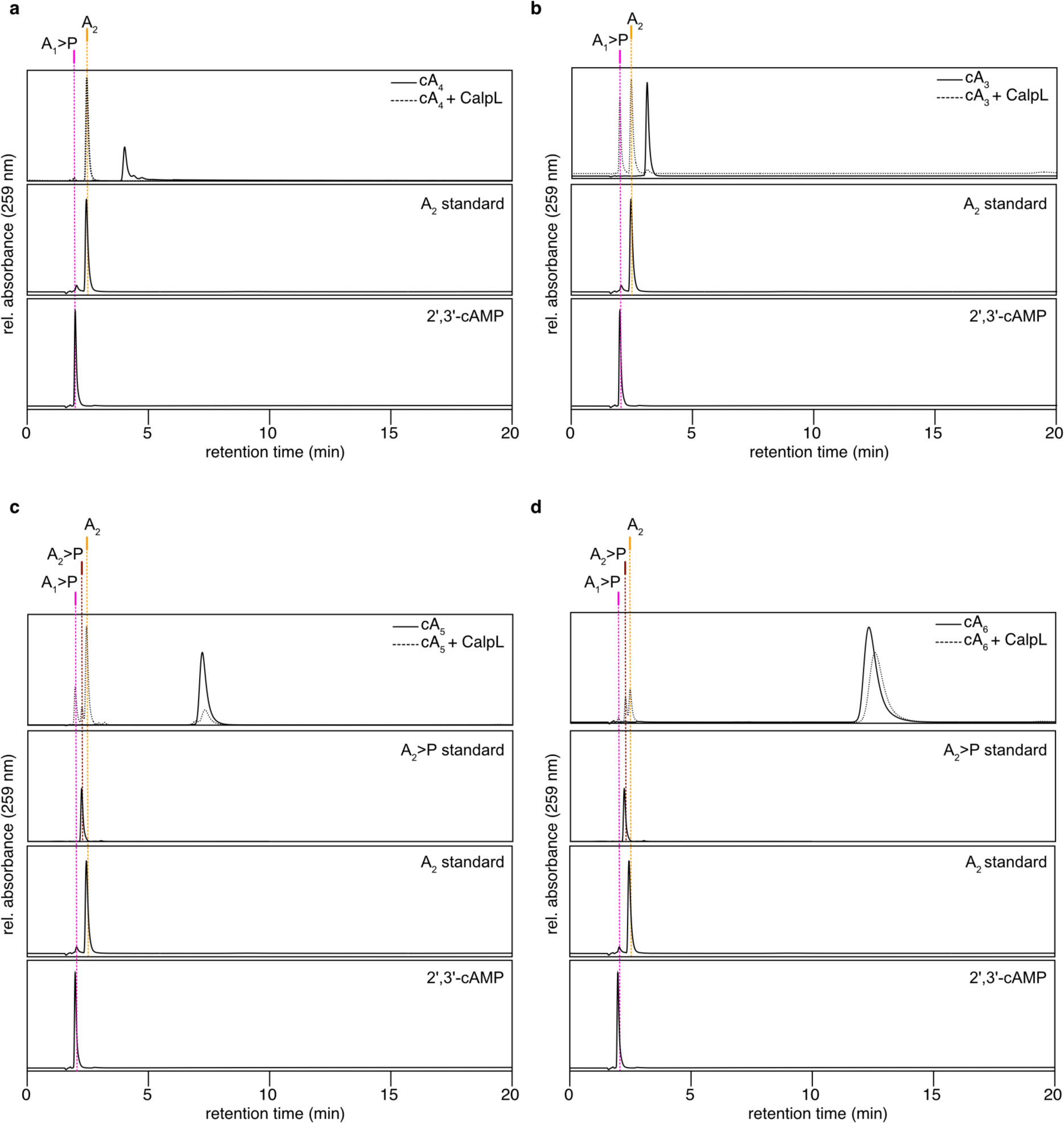
Identification of cOA degradation products by comparison to HPLC oligoadenylate standards. **a-d)** HPLC traces recorded at 259 nm showing the result of incubating different cOAs with (dashed lines) and without (solid lines) CalpL for 120 min. at 60 °C. The A_2_>P standard was generated by incubating MazF with the respective substrate for 60 min. at 37 °C. HPLC traces are representative of three replicates.

**Fig. S2.**
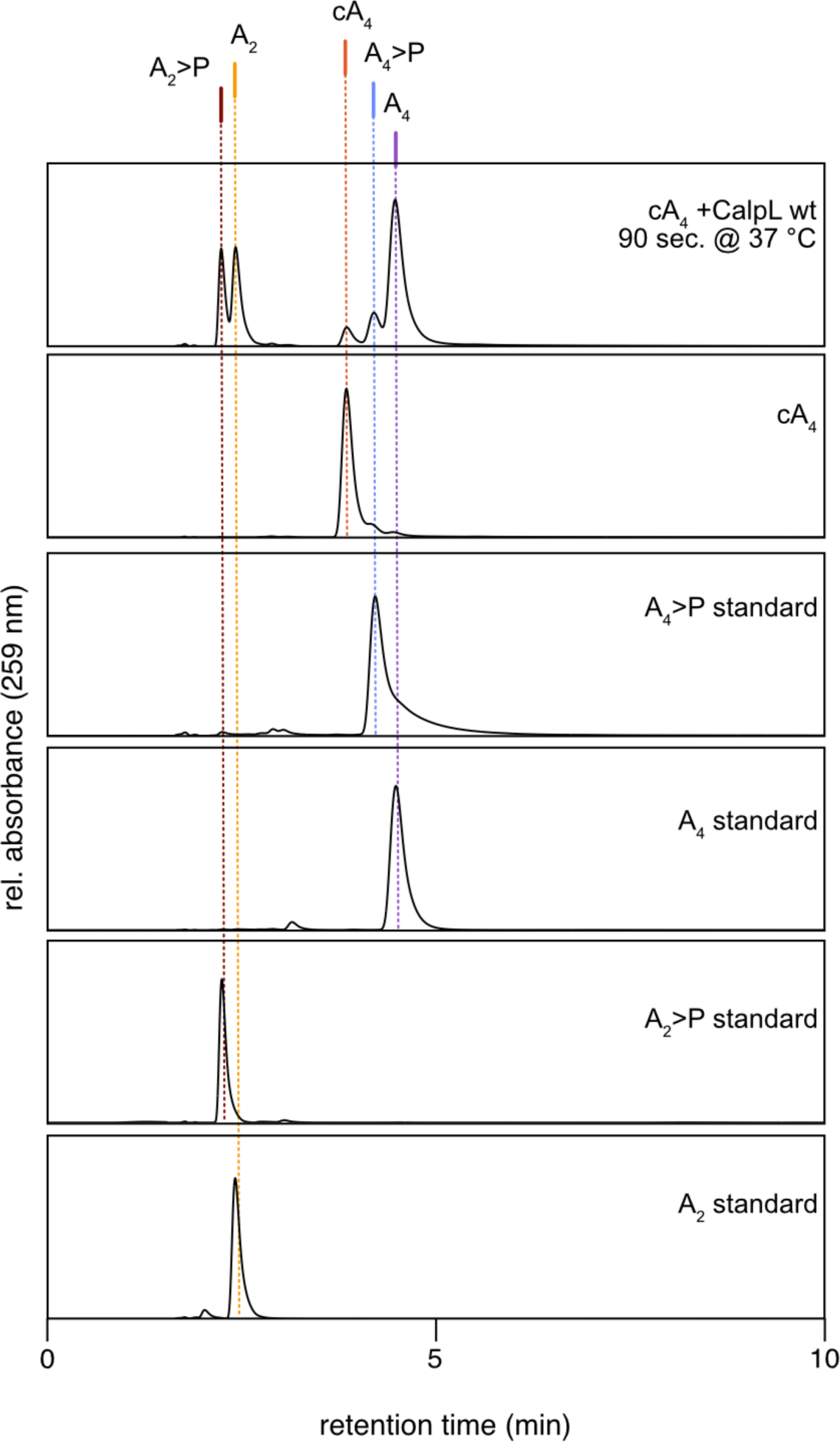
Identification of cA_4_ degradation products by comparison to HPLC oligoadenylate standards. HPLC traces recorded at 259 nm showing the reaction intermediates and product generated upon incubation of 30 µM cA_4_ with 3 µM CalpL at 37 °C for 90 sec. Standards containing cyclic phosphate groups were generated by incubating MazF with the respective substrate for 60 min. at 37 °C. HPLC traces are representative of three replicates.

**Fig. S3.**
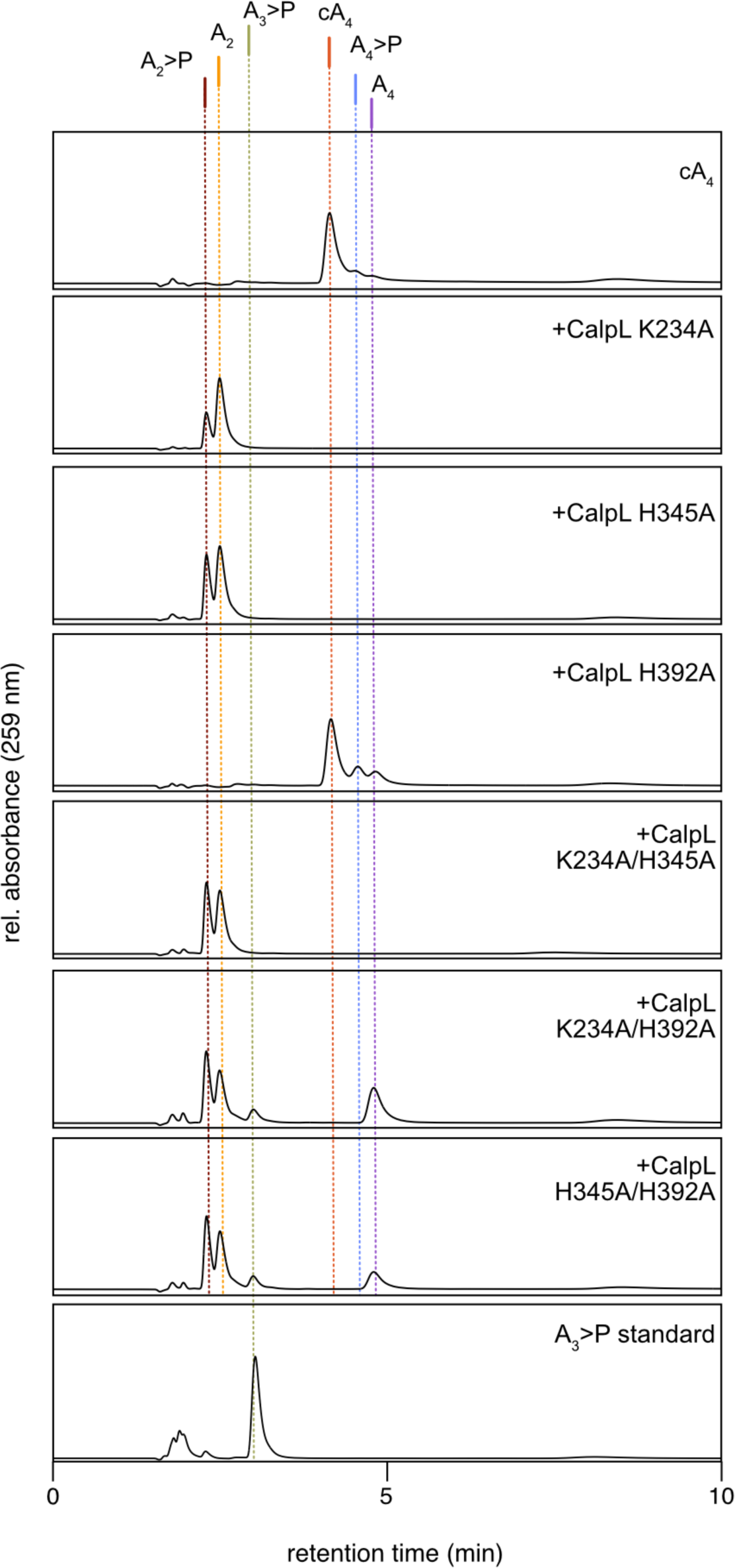
Nuclease reaction endpoints of SAVED topside double mutants. HPLC traces recorded at 259 nm showing the reaction intermediates and products generated upon incubation of 15 µM cA_4_ with 1.5 µM of the respective CalpL single or double mutant at 60 °C for 120 min. HPLC traces are representative of three replicates. The A_3_>P standard was generated by incubating MazF with the respective substrate for 60 min. at 37 °C.

**Fig. S4.**
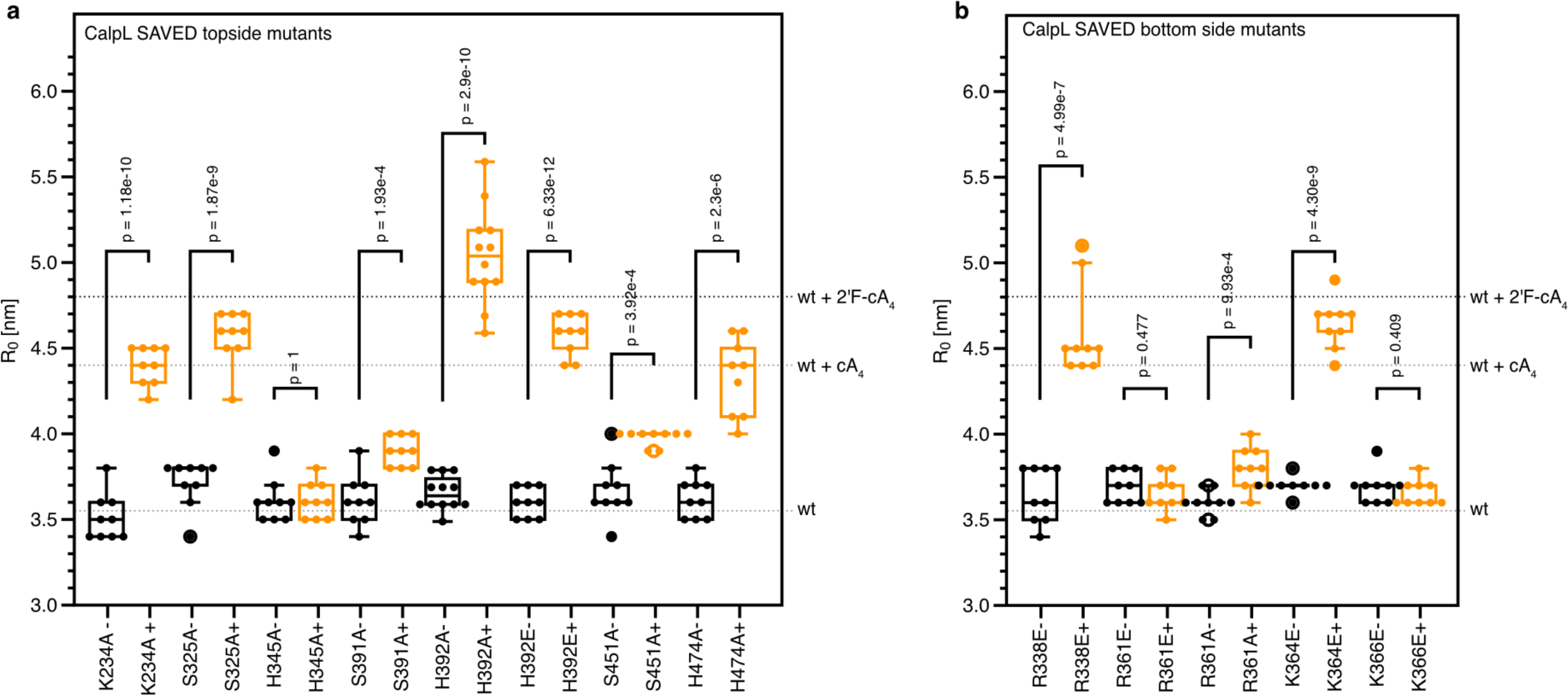
Hydrodynamic radii of SAVED topside and bottom side mutants. Hydrodynamic radii of SAVED topside (a) or bottom side (b) mutants determined in the presence (+; orange) and absence (-; black) of cA_4_ by dynamic light scattering (DLS). Mean hydrodynamic radii of wildtype CalpL in the presence of cA_4_ (grey traces) or 2’F-cA_4_ (black traces) are indicated. Measurements were performed at a sample concentration of 86 µM (5 mg/ml) and at a temperature of 20 °C. For each condition, three measurement cycles of each 20 single data acquisitions with acquisition times of 3 sec were recorded. Bars of DLS data display median and interquartile range; *p*-values for two-tailed t-tests are indicated.

**Fig. S5.**
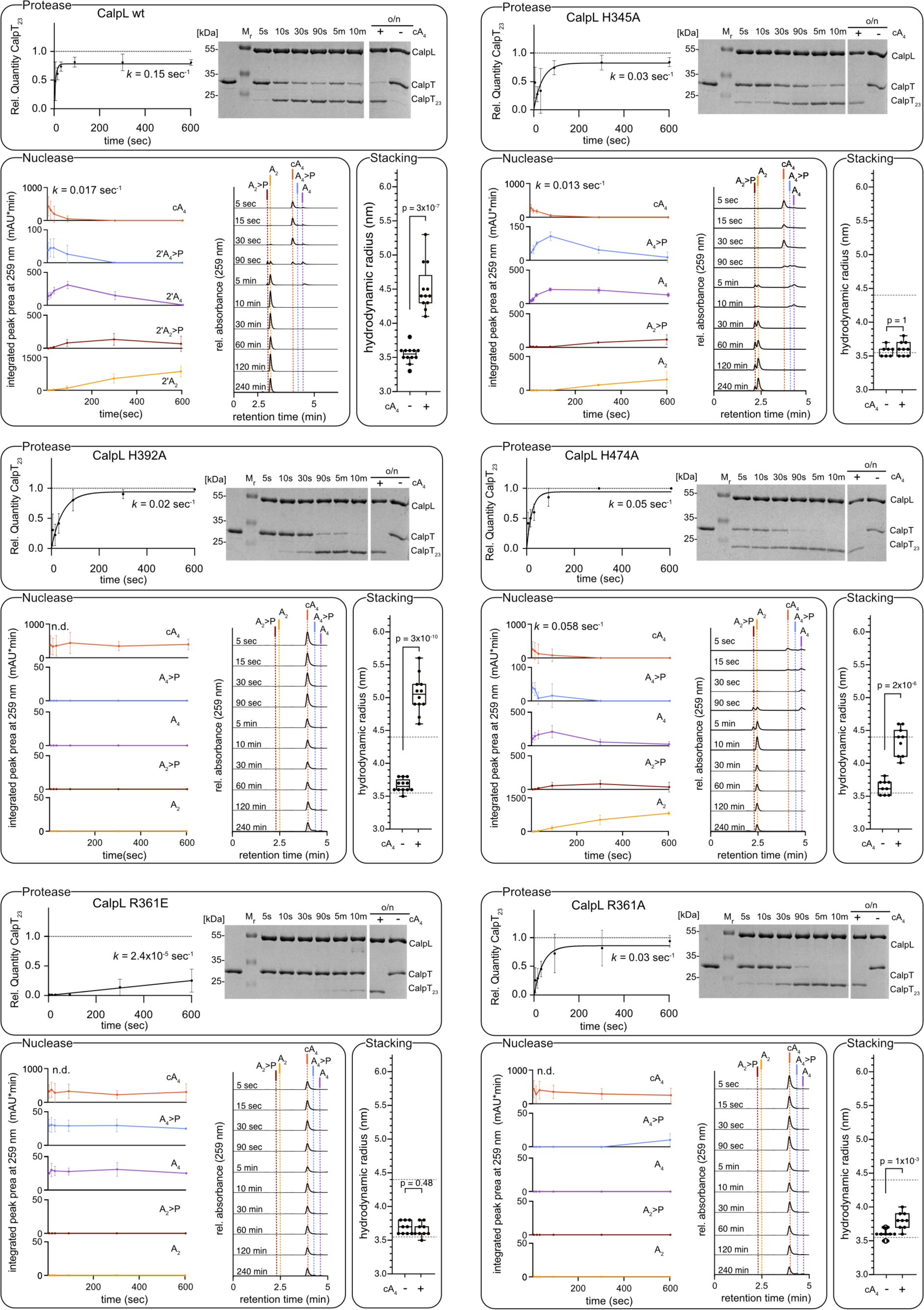
Interdependence of ring nuclease and protease activities of CalpL. Upper panels: Quantification of CalpT cleavage by incubating 3 µM of the respective CalpL mutant with 3 µM CalpT at 37 °C for the indicated time points. A Coomassie-stained SDS-PAGE gel used for quantification is included for reference. Left lower panels: Quantification of nuclease reaction intermediates and products generated upon incubation of 1.5 µM CalpL and 15 µM cA_4_ at 37 °C for the indicated time points. HPLC traces used for quantification by peak area integration at 259 nm are shown exemplary. Right lower panels: Hydrodynamic radii of CalpL determined at a sample concentration of 86 µM (5 mg/ml) in the presence or absence of cA_4_. Dashed lines mark the hydrodynamic radii of wildtype CalpL with (4.5 ± 0.35 nm) and without (3.5± 0.12 nm) cA_4_. All experiments were performed in triplicates. Bars of DLS data display median and interquartile range; *p*-values for two-tailed t-tests are indicated.

**Table S1.** Accurate mass determination from sample fractions collected after HPLC separation.

|  | signal<br>assignment | measured<br>*m/z* | calculated<br>*m/z* | elemental composition |
| --- | --- | --- | --- | --- |
| A<sub>4</sub> | [M-2H]<sup>2-</sup> | 666.1032 | 666.1030 | [C<sub>40</sub>H<sub>48</sub>N<sub>20</sub>O<sub>25</sub>P<sub>4</sub>]<sup>2-</sup> |
|  | [M-3H+Na]<sup>2-</sup> | 677.0893 | 677.0940 | [C<sub>40</sub>H<sub>47</sub>N<sub>20</sub>O<sub>25</sub>P<sub>4</sub>Na]<sup>2-</sup> |
| A<sub>4</sub>>P | [M-2H]<sup>2-</sup> | 657.0974 | 657.0978 | [C<sub>40</sub>H<sub>46</sub>N<sub>20</sub>O<sub>24</sub>P<sub>4</sub>]<sup>2-</sup> |
|  | [M-3H+Na]<sup>2-</sup> | 668.0843 | 668.0887 | [C<sub>40</sub>H<sub>45</sub>N<sub>20</sub>O<sub>24</sub>P<sub>4</sub>Na]<sup>2-</sup> |
| cA<sub>4</sub> | [M-2H]<sup>2-</sup> | 657.0951 | 657.0978 | [C<sub>40</sub>H<sub>46</sub>N<sub>20</sub>O<sub>24</sub>P<sub>4</sub>]<sup>2-</sup> |
|  | [M-3H+Na]<sup>2-</sup> | 668.0885 | 668.0887 | [C<sub>40</sub>H<sub>45</sub>N<sub>20</sub>O<sub>24</sub>P<sub>4</sub>Na]<sup>2-</sup> |
| A<sub>2</sub> | [M-H]<sup>-</sup> | 675.1096 | 675.1083 | [C<sub>20</sub>H<sub>25</sub>N<sub>10</sub>O<sub>13</sub>P<sub>2</sub>]<sup>-</sup> |
|  | [M-2H+Na]<sup>-</sup> | 697.0932 | 697.0903 | [C<sub>20</sub>H<sub>24</sub>N<sub>10</sub>O<sub>13</sub>P<sub>2</sub>Na]<sup>-</sup> |
| A<sub>2</sub>>P | [M-H]<sup>-</sup> | 657.1007 | 657.0978 | [C<sub>20</sub>H<sub>23</sub>N<sub>10</sub>O<sub>12</sub>P<sub>2</sub>]<sup>-</sup> |
|  | [M-2H+Na]<sup>-</sup> | 679.0869 | 679.0797 | [C<sub>20</sub>H<sub>22</sub>N<sub>10</sub>O<sub>12</sub>P<sub>2</sub>Na]<sup>-</sup> |

## Supplementary Material: Sequence alignment underlying Fig. 4 and Fig. 5

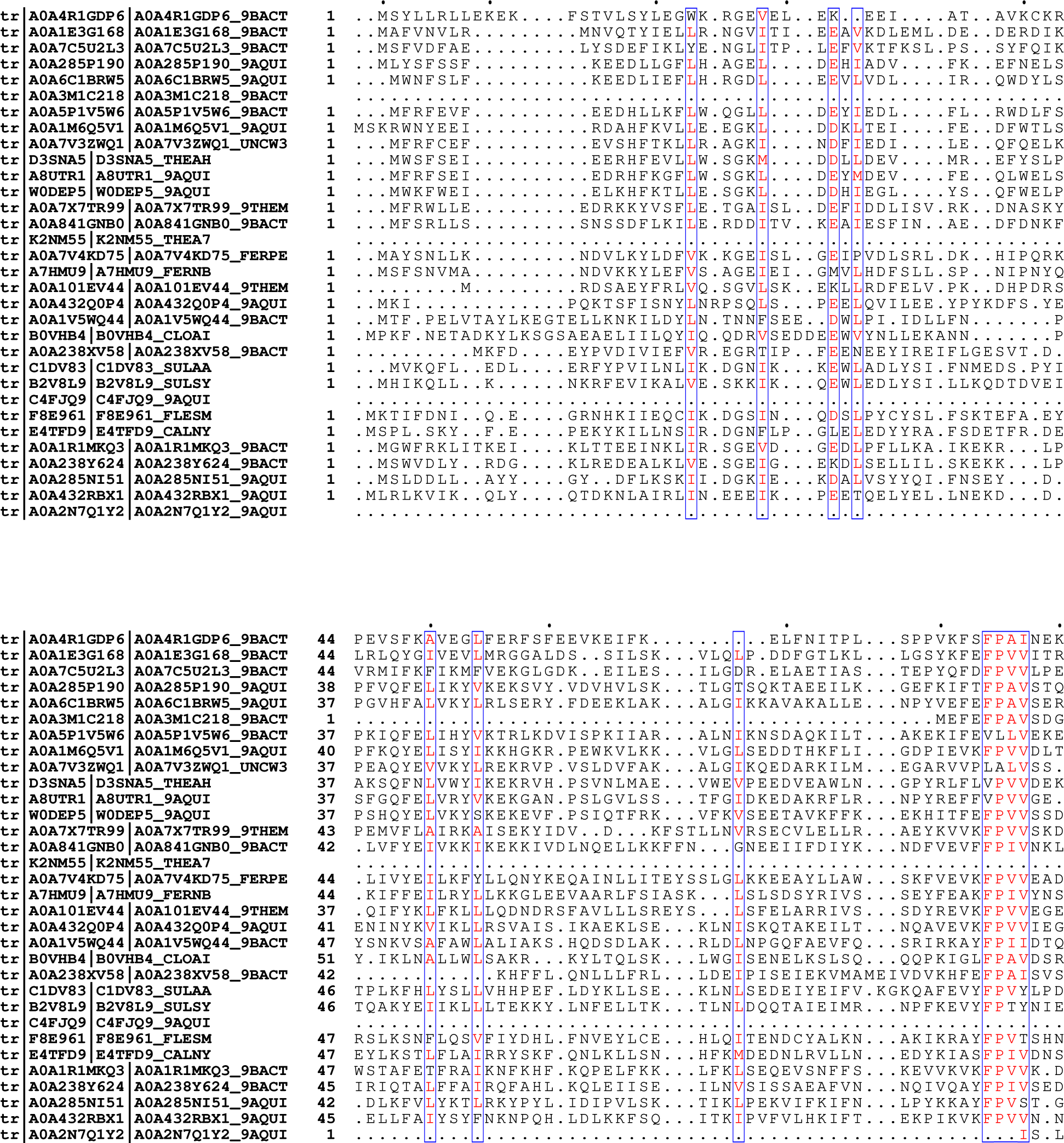
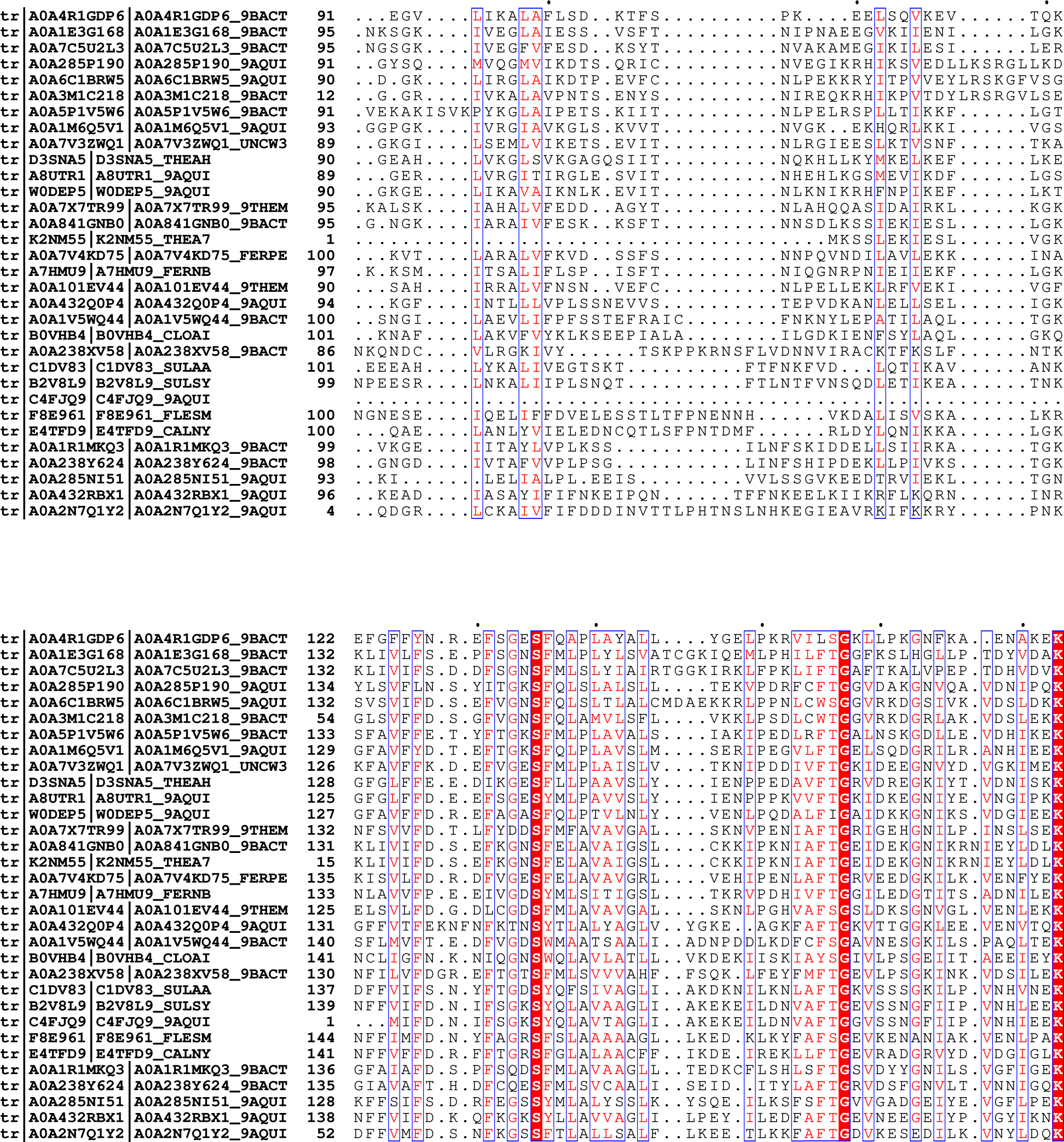
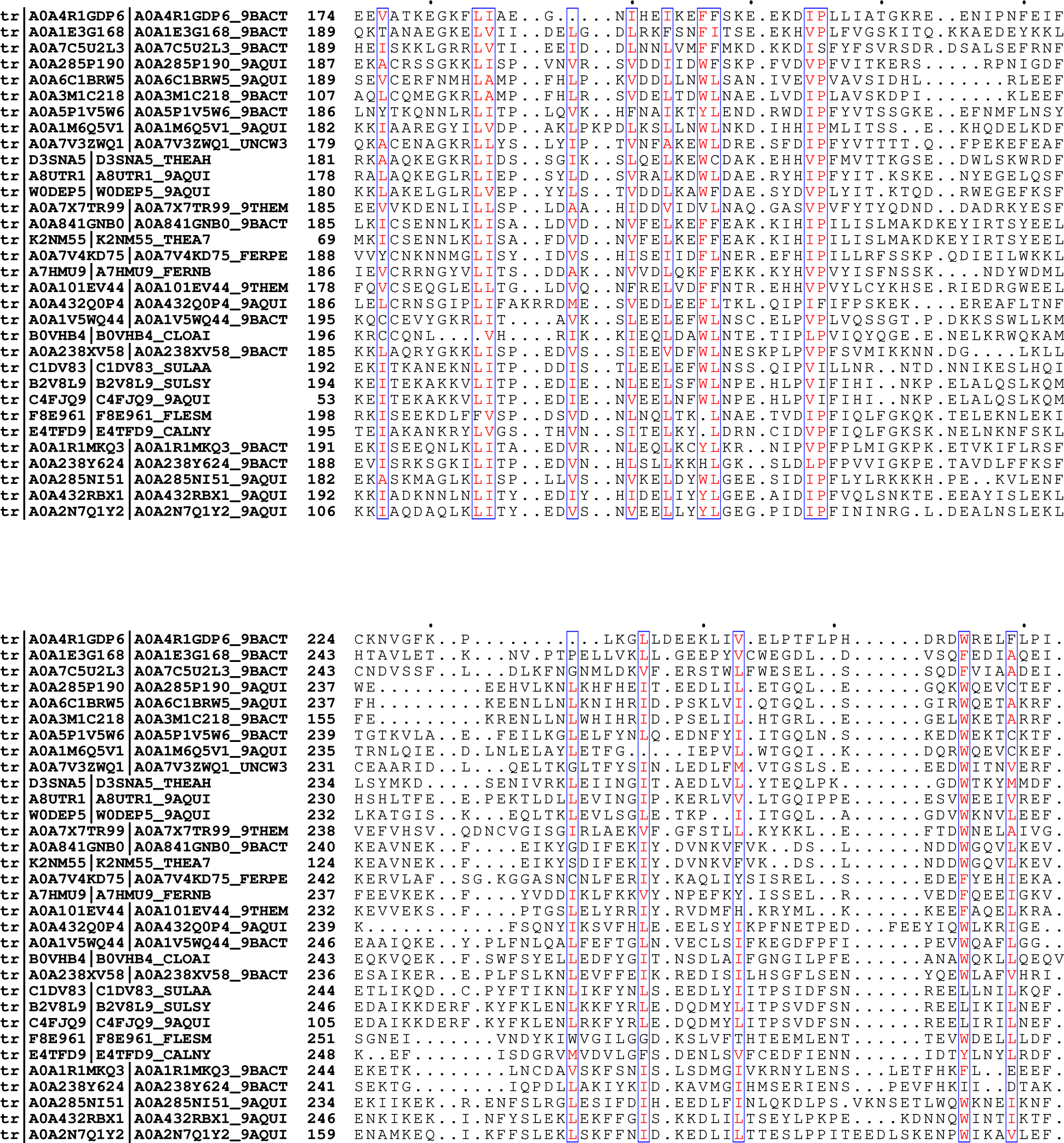
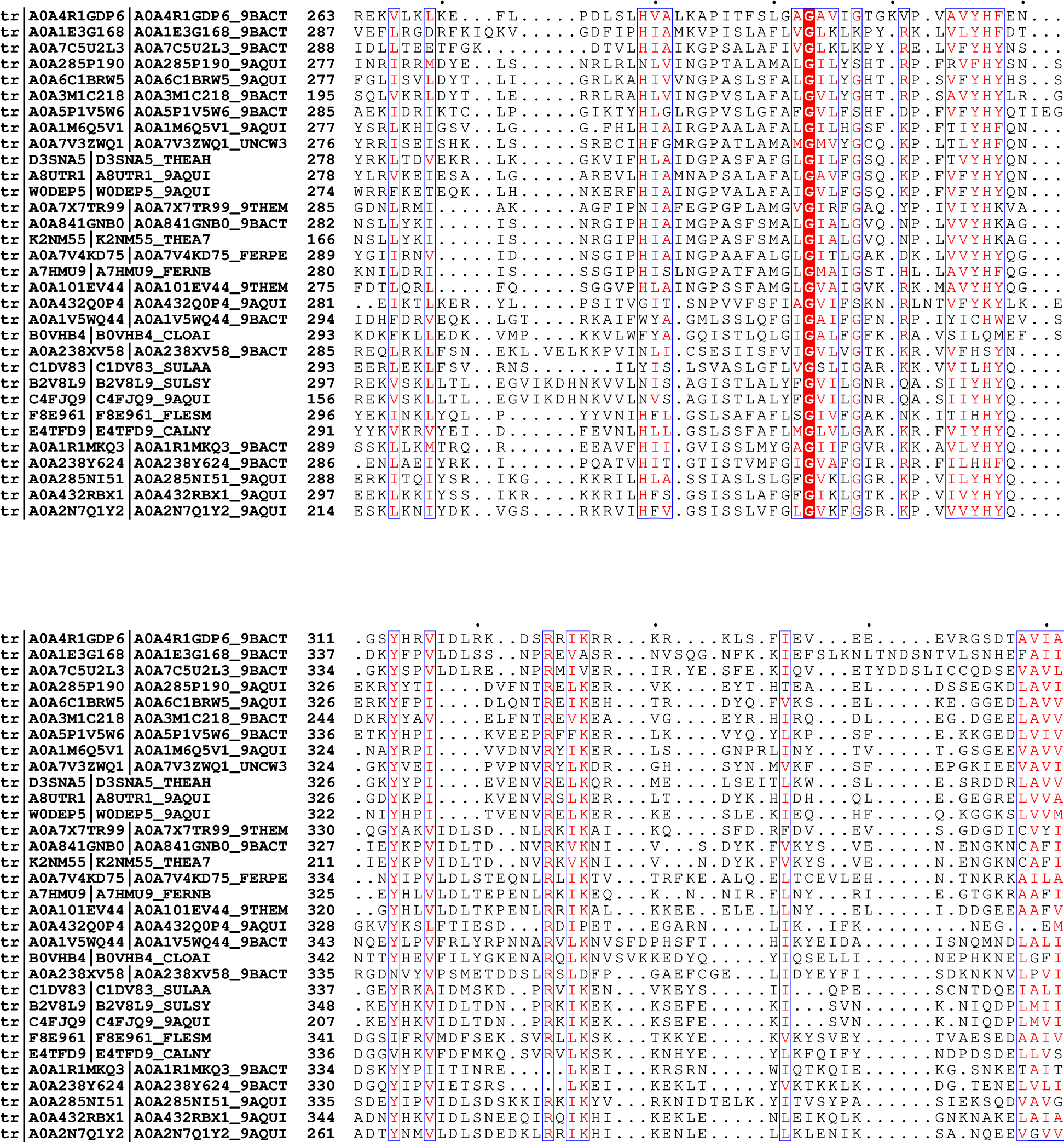
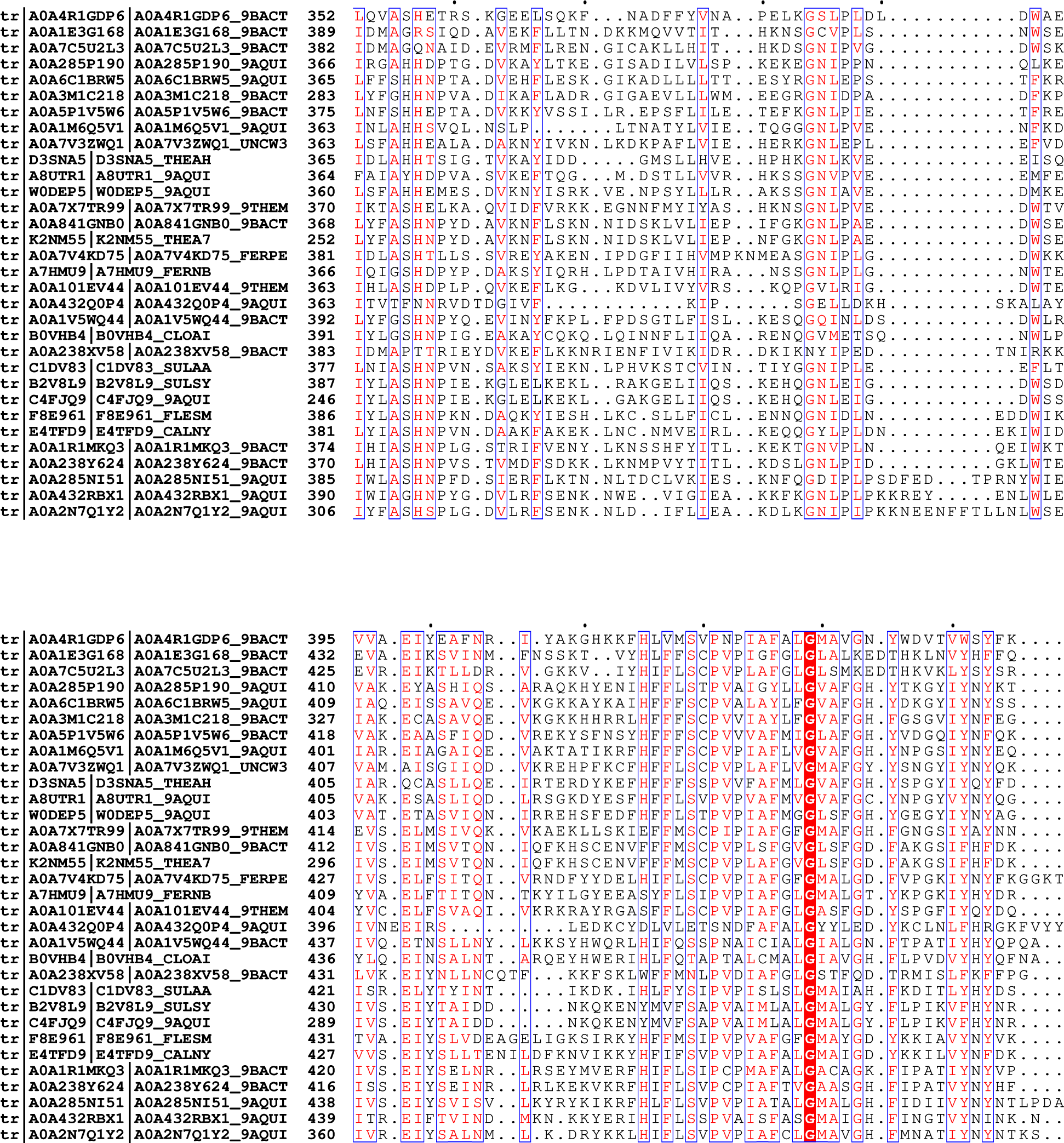
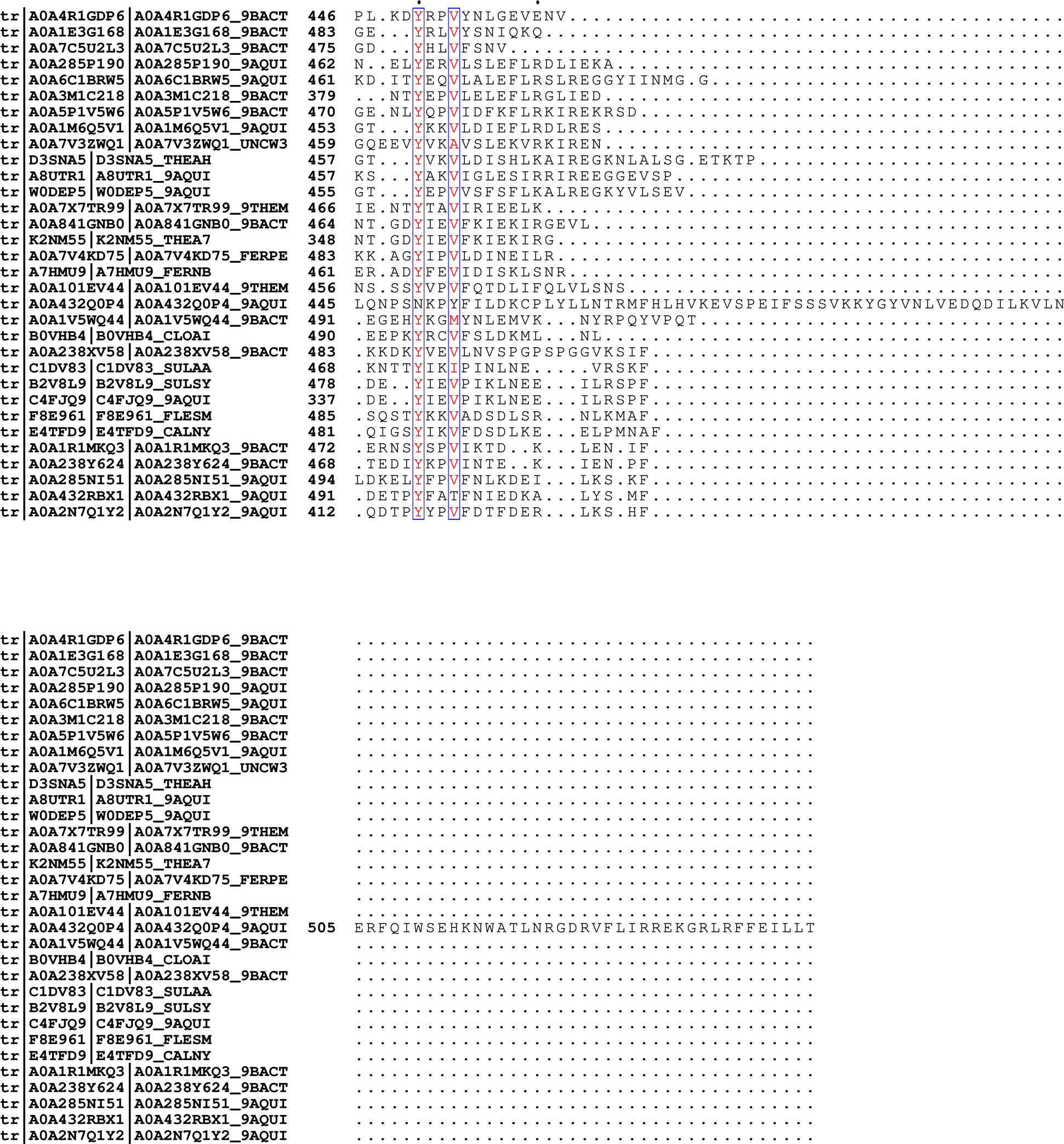

## Notes

### Competing Interest Statement

The authors have declared no competing interest.

